# First analysis of behavioural responses of humpback whales (*Megaptera novaeangliae*) to two acoustic alarms in a northern feeding ground off Iceland

**DOI:** 10.1101/741553

**Authors:** Charla J. Basran, Benno Woelfing, Charlotte Neumann, Marianne H. Rasmussen

**Affiliations:** Department of Life and Environmental Sciences, University of Iceland, Reykjavik, Iceland; Húsavík Research Centre, University of Iceland, Húsavík, Iceland; Institute for Terrestrial and Aquatic Wildlife Research, University of Veterinary Medicine, Hannover, Germany; University of Southern Denmark, Odense, Denmark

## Abstract

Mitigating cetacean entanglement in fishing industries is of global interest. Strategies include the use of acoustic alarms to warn whales of fishing gear. For baleen whales, responses to acoustic alarms are poorly understood. This behavioural response study compared the behaviour of humpback whales (*Megaptera novaeangliae*) in their feeding grounds off Iceland prior to, during, and after exposure to a low-frequency whale pinger (Future Oceans) and a high-frequency seal scarer (Lofitech ltd.). Linear mixed effects models and binary generalized linear mixed effects models were used to analyze the effect of the alarms on surface feeding, swimming speed, breathing rate, directness and dive time. We observed a significant decrease in surface feeding and a significant increase in swimming speed during exposure to the whale pinger. Changes in dive time between the phases of a trial differed significantly between individuals indicating that responses may depend on individual or behavioural state. We did not find any significant reactions in response to the seal scarer. In addition to the experimental exposures, a trial of whale pingers on a capelin purse seine net was conducted. Results from this trial showed that whales entered the net from the bottom while the pingers were attached at the top, but the encircled whales were able to locate an opening free of pingers and escape without damaging the net. Our results suggest that whale pingers may be a useful entanglement mitigation tool in humpback whale feeding grounds given that a reduction in feeding around nets likely reduces the risk of whales swimming through them. Pingers may also minimize net damage if whales are encircled by aiding the whales in finding their way out. However, given the uncertain long-term consequences of the behavioural changes reported here, whale pingers are most advisable for short-term use in conjunction with other entanglement mitigation measures.

## Introduction

There is global concern over marine mammal bycatch and entanglement in fishing gear (ie. animals becoming incidentally caught in gear and drowning, or escaping, sometimes with gear attached to their body and/or with injuries). Documented impacts of entanglement on cetaceans include injury [1, 2, 3], exhaustion of energy budgets [4], emaciation and drowning [5, 2]. These impacts at the individual level can lead to increased mortality rates at the population level [6, 7]. Entanglement is known to occur involving many different types of fishing gear [8] and is likely to affect most cetacean species [9]. Apart from impacts on cetacean individuals and populations, entanglement also leads to financial losses to the fishing industry due to loss-of-catch, gear damage or loss and downtime for repairs [10, 11]. This can be a particularly serious issue in fisheries experiencing large whale, such as humpback whale (*Megaptera novaeangliae*), entanglements.

Technologies have been developed with the intent to mitigate marine mammal entanglement. One such technology is acoustic alarms known as “pingers”. These devices can be to attached to fishing gear and emit a tone underwater within the hearing range of target marine mammals [12]. The devices serve to “illuminate” the gear with sound and warn the animals of its presence to encourage them to avoid it. Alternatively, the pingers may simply serve as an annoying, unnatural sound that the animals want to avoid [13]. Since large whales in particular can often escape from or carry away entangling gear, it has been suggested that whales can learn to associate nets and pingers with danger [14]. For cetaceans, specific pinger acoustic alarms have been developed for porpoises, dolphins and beaked whales (odontocetes) as well as baleen whales (mysticetes) with varying degrees of success. The high-frequency porpoise, dolphin and beaked whale pingers have been shown to reduce bycatch of several species [eg. 15, 16, 17, 18], though on the other hand, some studies have found there to be no change or even increased bycatch of some odontocete species with the use of pingers [eg. 19, 20]. Whale pinger and low-frequency acoustic alarm sound experiments have been conducted on baleen whales, including North-Atlantic right whales (*Eubalaena glacialis*) [21], minke whales (*Balaenoptera acutorostrata*) [22], grey whales (*Eschrichtius robustus*) [23], and humpback whales [24, 25, 26, 27, 22]. Results from these experiments were also highly variable. North-Atlantic right whales showed a strong response to alerting sounds [28], and both minke whales and humpback whales also responded during testing of whale alarm prototypes [22]. Humpback whales were also less likely to collide with cod-traps fitted with alarms in Canada [27] and were found to respond to “tone stimuli” within their hearing range in Australia [29]. By contrast, grey whales did not appear to respond to acoustic deterrent sounds though results were inconclusive [23] and the majority of recent research conducted on humpback whales in Australia has concluded there is no clear response to the modern whale pinger alarms, some of which are now sold commercially [24, 25, 26]. Despite this, anecdotal reports do claim that some industries have had lower incidence of humpback whale entanglement with the use of the commercial whale pingers [30, 31].

In addition to the pinger acoustic alarms, acoustic deterrent devices (ADDs) have also been developed. Primarily used in the aquaculture industry to ward off seals, these devices produce a loud, high-frequency sound designed to scare away the animals [32]. Though not designed originally for use to deter cetaceans, it has been observed that at least some cetacean species react to the loud sound produced by such a device [33, 34]. Testing of ADDs on cetaceans has found that odontocetes including harbour porpoises [34], orcas [35] and Pacific white-sided dolphins [36] are deterred by the devices. The only testing of such a device on baleen whales was conducted on minke whales in Iceland, and results showed that they too were deterred from the area with an active ADD [33].

Humpback whales are one of the most common cetaceans that frequent the waters off Iceland in the North Atlantic primarily during their feeding season from spring through autumn [37], though some sightings are also recorded in the winter months [38]. The summer-time Icelandic humpback whale population is estimated to be approximately 12,000 individuals [37] with the highest concentration found off the north/northeast coast (Pike et al. 2019 submitted). It is thought that the humpback whales’ diet in Iceland consists of 60% fish species [39]. The hearing capabilities of humpback whales has been modelled to show that they have hearing sensitivity between 700 Hz and 10 kHz, with maximum sensitivity between 2-6 kHz [40]. It has been suggested that hearing is likely the most important sense for baleen whales to orient themselves in their environment [41] and large baleen whales, like the humpback, may have trouble acoustically detecting fishing gear in the water.

Commercial fishing is one of the largest industries in Iceland, with 1588 commercial vessels registered in 2018 [42]. The fishing methods used in Icelandic waters are long-line/hand-line, gillnet, trawl, and purse seine [43]. In addition, there are also mussel, oyster, and fish farming operations in coastal Icelandic waters [44, Kristján Phillips pers. comm. 2019, 45]. At least one-quarter of the coastal Icelandic humpback whale population is estimated to have been entangled in fishing gear at least once [46], and virtually all of the fishing methods in the country have reported issues with humpback whales swimming through, and sometimes becoming entangled in, the gear in the water [47, 44]. This has caused gear damage or loss, as well as injury or death to the whales in some cases [46, 47, 48, 49].

Currently there are no mitigation methods or regulations in place for minimizing whale entanglement in fishing gear in Iceland, despite growing concern in the local fishing industries. This study conducted the first analysis of behavioural response of free-ranging humpback whales in their northern feeding grounds off the coast of northeast Iceland to the whale pinger acoustic alarm (Future Oceans) and the seal scarer acoustic deterrent device (Lofitech AS ltd.). In addition to the experimental exposure of whales to the acoustic alarms, this study conducted the first trial of the whale pingers in the capelin purse seine fishery in Iceland. Results from this study help to decide if acoustic alarms are likely to effectively mitigate humpback whale entanglement in their feeding grounds and shed light on possible adverse effects of the alarms on natural humpback whale behaviour.

## Materials and Methods

### Study area

Trials of the whale pinger acoustic alarm took place in two locations in Northeast Iceland: Skjálfandi Bay and Eyjafjörður (Fig 1). Skjálfandi (66°05’N17° 33’W) is an approximately 11,000 km^2^ bay well known for predictable humpback whale sightings from spring through autumn during the feeding season. The bay harbours the fishing-turned-whale watching town of Húsavík on the southeast shore [50, 51, 52, A. Gíslason unpubl. data]. Eyjafjörður (65°50’N18° 07’W) is an approximately 440km^2^ narrow fjord (S. Jónsson unpubl. data) located approximately 80km west of Skjálfandi Bay. Like Skjálfandi, Eyjafjörður is also well known for humpback whale sightings and harbours fishing and whale watching in the city of Akureyri as well as the towns of Dalvík, Hauganes and Hjalteyri. Trials of the seal scarer acoustic alarm took place only in Skjálfandi Bay.

**Fig 1.**
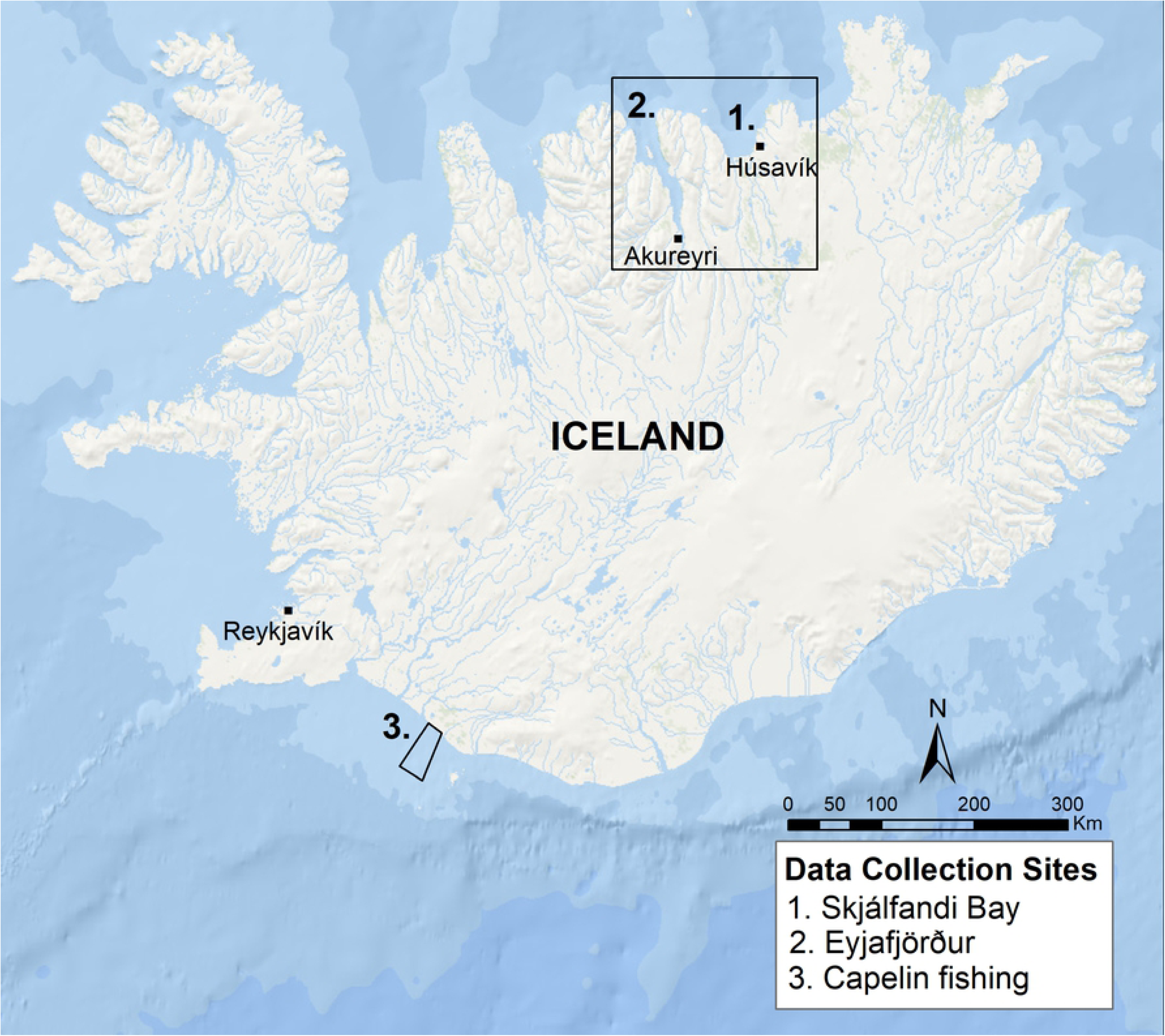
Map showing the locations of humpback whale behavioural response trials (1. and 2.) using the whale pinger and/or seal scarer, and location where capelin fishing with a purse seine net equipped with the whale pingers took place during onboard observation (3.).

The practical trial of the whale pinger took place in collaboration with a capelin purse seine vessel based in Neskaupstaður in East Iceland. The boat fished for capelin off South Iceland (Fig 1).

### Acoustic alarms

Two acoustic alarms were used in the present study. First was the 2016 version of the Future Oceans whale pinger. This device operates on a single 3.6V lithium battery and activates automatically when in contact with saltwater. When active, the alarm produces a 145 decibel re 1µPa tone at 3kHz for 300 ms at 5 sec intervals (Future Oceans) (Fig 2). The second alarm was the Lofitech AS ltd. seal scarer ADD composed of a control box with a 25m long cable with a transducer unit at the end which produces the sound. This control box is powered by a 12V marine battery onboard the boat. When active, the alarm produces a 191 decibel re 1µPa sound between 10-20 kHz for 500ms at random intervals of 5-60 sec (Fig 3). A calibration of both acoustic alarm devices was conducted in a harbour to confirm the manufacturers specifications. Each device was lowered 5m into the water and recorded by a Reson 4032 hydrophone connected to an Etec amplifier with the sound signal recording to a Microtrack recorder. The whale pinger was recorded at distances of 1, 5, 10 and 20m from the hydrophone, while the seal scarer was recorded at 20, 30 and 40m. The recorded signal from the alarms was compared with a 153 dB rms calibration signal recorded using a calibrator with an adapter for the 4032 Reson hydrophone.

**Fig 2.**
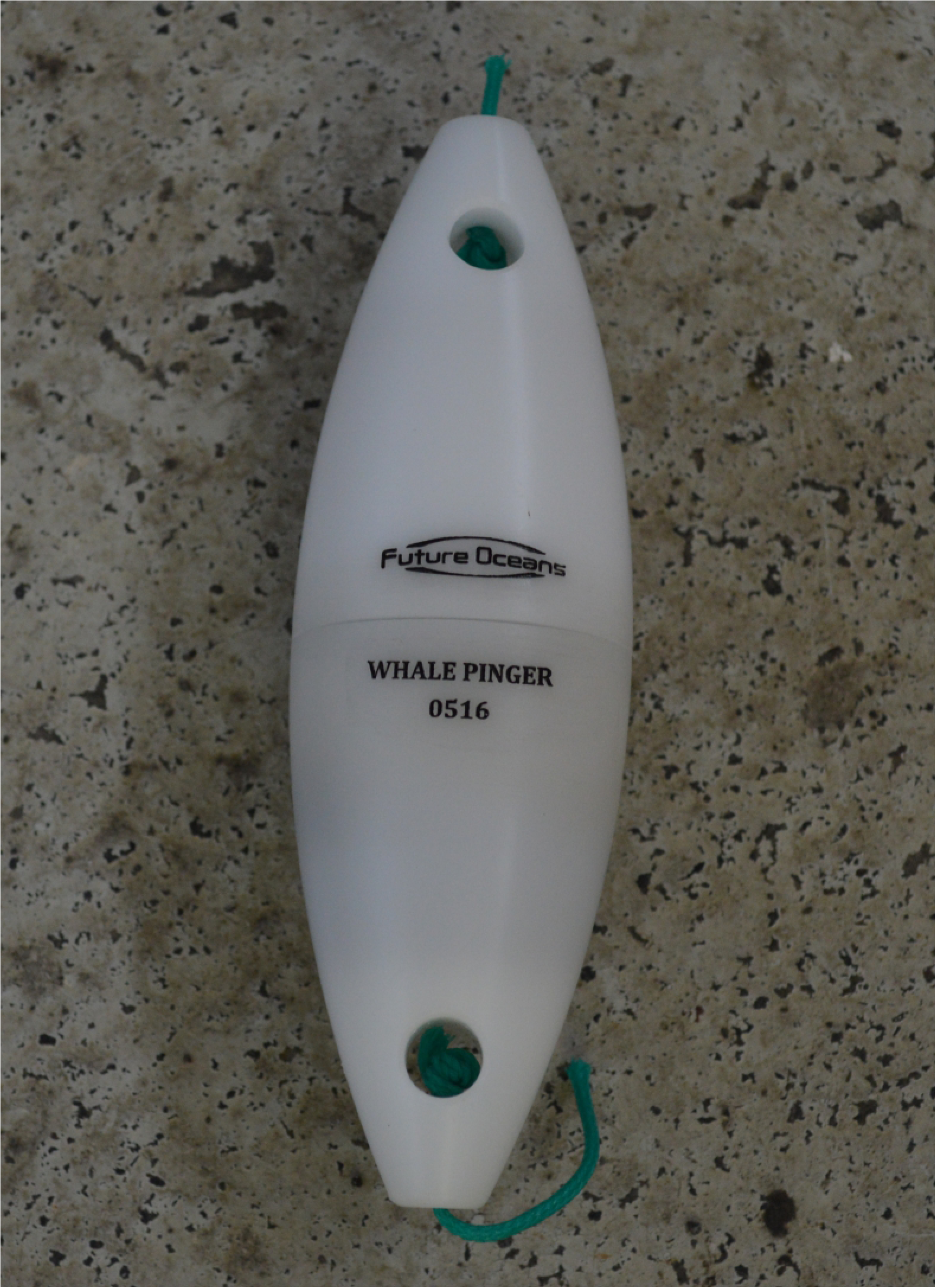
Photograph showing the whale pinger alarm.

**Fig 3.**
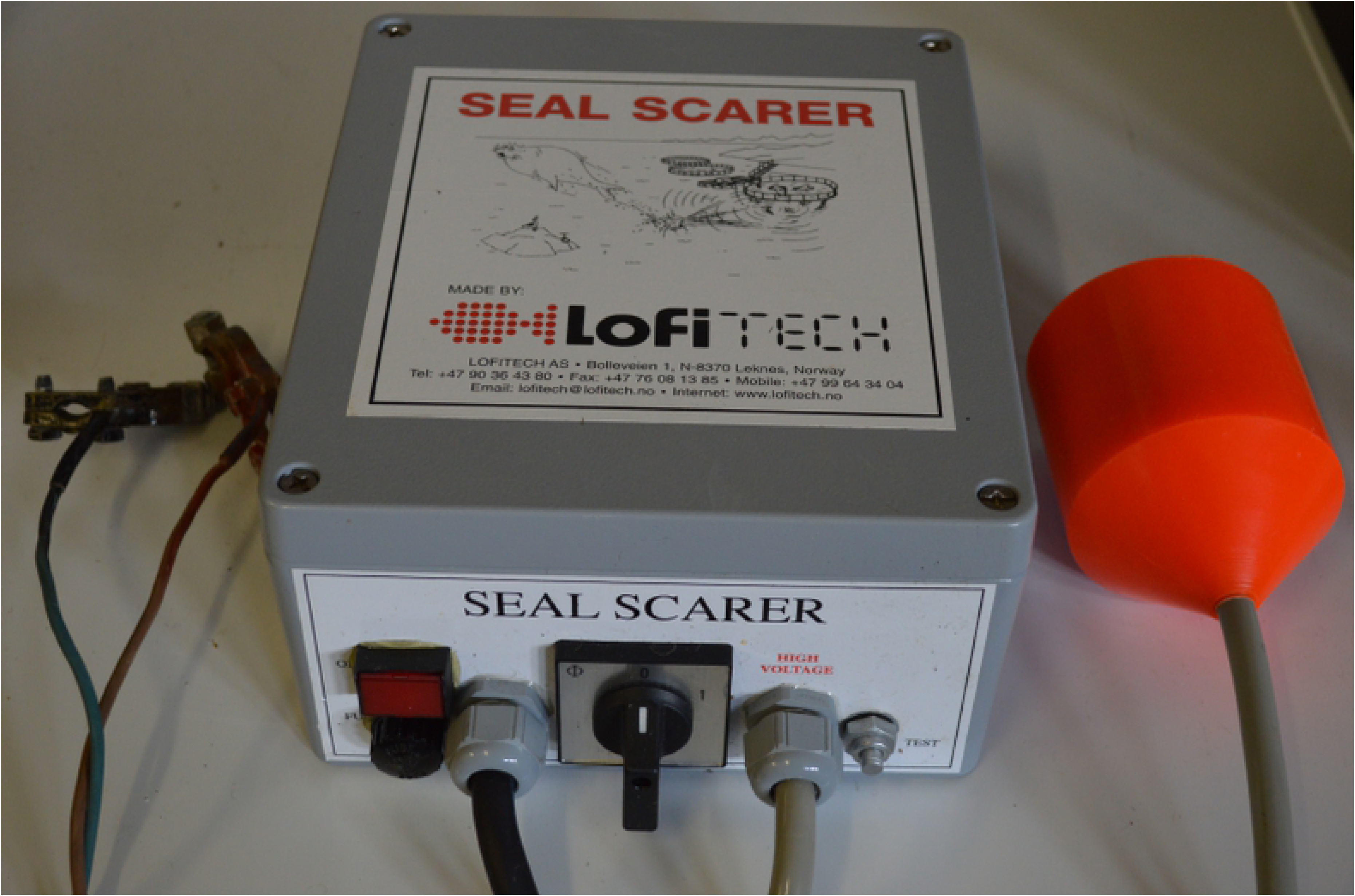
Photograph showing the seal scarer alarm.

The emitted sound from the whale pinger had an actual source level of 137 dB re 1µPa (rms) recorded at a distance of 1 m. The levels at 5, 10 and 20 m were 137 dB, 140 dB and 144 dB re 1µPa (rms) respectively using spherical spreading as transmission loss. Based on previous modelling of the pinger sound, humpback whales are expected to detect the sound at a distance of at least 500m from the source [24, 26]. The seal scarer had an actual source level of 189-198 dB re 1µPa (rms) measured at a distance of 20, 30 and 40 m using spherical spreading (of 20 Log R) as transmission loss.

### Experimental exposure to acoustic alarms

Experimental exposures of humpback whales to acoustic alarms were conducted in Skjálfandi Bay in June, July and October 2017, and in June and October 2018. In Eyjafjörður, trials were conducted in May and July 2018. A private boat was used for each trial with a captain and 3-4 researchers and students onboard. Data collected during the trials were recorded with the Logger 2010 computer program (IFAW) and a video recording of each trial was taken using a hand-held video camera (Sony HDR-CX160E handycam). Logger 2010 recorded time, GPS position of the boat, heading of the boat, and any comments that were entered by the student recorder. Trials were attempted when the sea state was considered 3 or less on the Beaufort scale. During a behavioural trial, an individual focal humpback whale was chosen based on the criteria that it was swimming alone and that there were no whale watching boats observing the animal. Photo-identification images of the individual were taken of the unique pattern on the underneath side of the fluke and of the dorsal fin. This was to ensure each individual whale was not exposed to the same device more than once within the same year, to avoid possible habituation to the alarm sound. When photo-identification was complete, the pre-exposure phase (*PrE*) began with the boat following the focal whale from a distance of approximately 100m for 30 mins to obtain a baseline of behaviour of the individual. The 100m distance complies with whale watching criteria set forth in many countries around the world to minimize disturbance to the animal [53] while still being within range to collect all necessary data. Each breath the whale took was recorded as “up” and each terminal dive was recorded as “dive” in Logger 2010. Other information was also noted, including if the whale dove with or without raising the fluke, if the whale appeared to be feeding, and if there were other whales in the area. Furthermore, one researcher used an angle-board and rangefinder to obtain the angle to the whale in relation to the boat and the distance to the whale, and this data was also recorded into Logger 2010. If the distance could not be obtained from the rangefinder, one researcher estimated the distance to the whale when it took a terminal dive. The angle-board, rangefinder, and distance estimation were always done by the same researcher (C.J.B) for consistency. Once the *PrE* phase was complete, the boat was positioned beside where the focal whale was seen taking its last terminal dive and the engine was turned off. To begin the 15 min exposure phase (*E*), the whale pinger or the seal scarer was placed off the side of the boat into the water at 5m depth, attached to a rope and buoy similar to Harcourt et al [26]. The breaths, dives, angles, and distances of the focal whale were then recorded in Logger 2010 in the same manner as in *PrE* phase. After the 15 min *E* phase ended, the alarm device was removed from the water and the boat was positioned approximately 100m from the focal whale to follow it and record the same data for an additional 30 mins for the post-exposure phase (*PoE*).

### Behavioural variables

#### Feeding

The number of surface feeding events was determined by watching the video footage of each phase of each behavioural trial. For each surfacing of the focal whale, surface feeding behaviour was categorized as yes (Y), no (N), or not able to determine (NA). Feeding behaviour was recognized by observing surface lunging behaviour or expanded throat pleats indicating the whale had a full mouth (Fig 4). A surfacing was also categorized as Y if researchers audibly indicated the whale was feeding in the video even though the surfacing was not visible in the footage.

**Fig 4.**
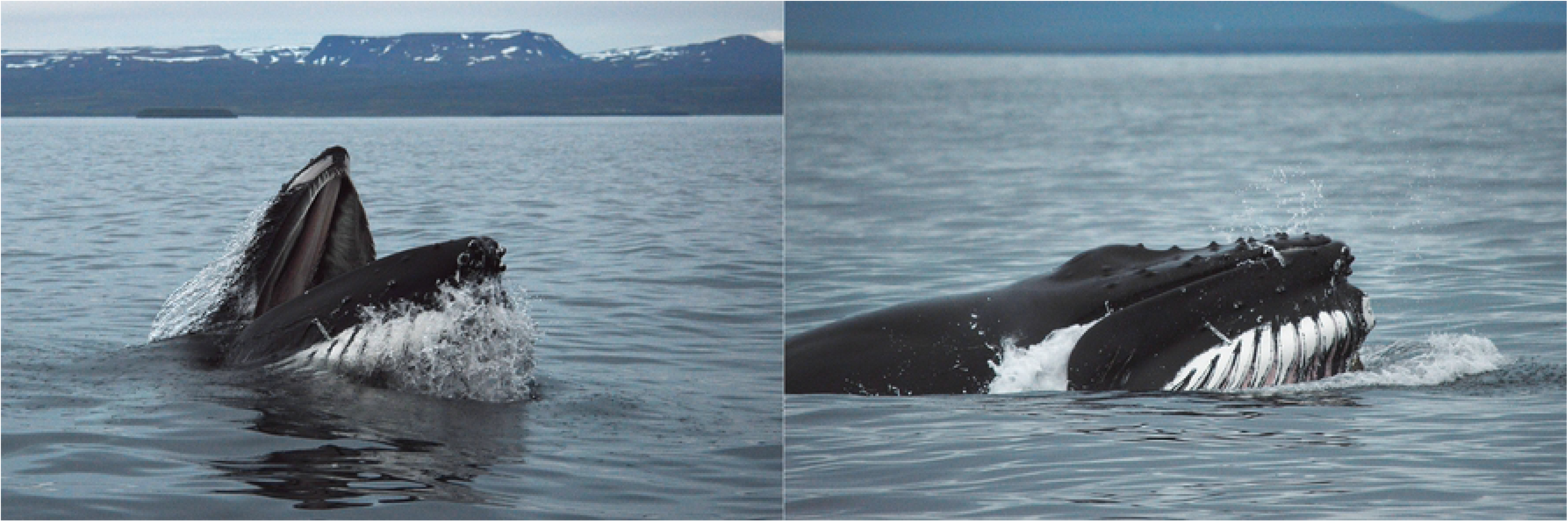
Photographs showing lunge-feeding behaviour and expanded throat pleats used to determine if the focal whale was surface feeding in the analysis of the videos.

#### Swimming speed

The swimming speed of the focal whale was calculated for each phase of each behavioural trial, when enough data was available. Speed was calculated from each terminal dive to the next terminal dive (and therefore included distance information from when the focal whale was diving and was at the surface).

#### Breathing rate and dive time

For each phase of each behavioural trial, the breathing rate of the focal whale was calculated as breaths per minute for each surface interval (the time between diving). The time of each dive in seconds in each phase of each trial was also calculated from the time stamps of “dive” and the following “up” recorded in Logger 2010.

#### Directness index

A directness index (DI) from 0-100, indicating the directness of the swimming pattern of the focal whale, was calculated for each phase of each behavioural trial, when enough data was available. Firstly, the coordinate position of the whale at each terminal dive was calculated. Then, the DI was calculated as the distance between the two end points of the track divided by the sum of the distances between all the points in the track, and the result multiplied by 100. A DI of 0 indicates swimming in a complete circle, while a DI of 100 indicates swimming in a straight line.

### Analysis of behavioural response variables

We tested the effect of exposure to both acoustic alarms (whale pinger and seal scarer) on four response variables: speed, breathing rate, directness and dive time using linear mixed effects models. Separate models were set up for each acoustic alarm and each response variable. The phase of the trial (*PrE*, *E*, *PoE*) was the only fixed effect predictor variable. To account for the repeated measures within individual whales, trial-ID was included as a random intercept term in all models. Plots of residual versus fitted values revealed that speed and breathing rate needed to be log-transformed to satisfy the modeling assumption of homogeneity of variances. Plots of the autocorrelation function of the residuals revealed significant temporal autocorrelation in the models for ln(speed), ln(breathing rate) and dive time. Auto-regressive correlation structures of order 1 were specified in the models for these response variables. Inspection of the plots of the autocorrelation functions verified that this successfully accounted for the observed autocorrelation.

Since previous findings suggested individual response to sound can depend on behavioural state [54], individual-specific response variation was incorporated into our models by introducing random slopes for the predictor phase for all response variables. We tested if random intercept and slope models fitted the data better than pure random intercept models. As recommended by Zuur et al. [55] selection of the random effects structure was done prior to selection of the fixed effects structure and was based on a likelihood ratio test comparing the pure random intercept model with the random intercept and slope model. Subsequently, likelihood ratio tests were used to select the optimal fixed effects structure, i.e. to compare models with phase as fixed effect to pure intercept models. For models in which phase had a significant effect, a posthoc pairwise comparison with Bonferroni correction was used to infer between which phases significant changes of the response variable occurred. These statistical analyses were performed using the libraries nlme [56] and multcomp [57] in the statistical software R (R Foundation for Statistical Computing).

Surface feeding behaviour was recorded as a binary variable and thus could not be modelled by linear mixed effects models. We fitted a binary generalized linear mixed effects model using the function glmer in the lme4-package [58]. Model specification and selection was analogous to the protocol described for the linear mixed effects models except for the specification of the autocorrelation structure. Since the glmer-function does not allow for the specification of temporal correlation structures, the feeding behaviour at the previous surfacing event (lag1_feeding) was included as a fixed effect to account for temporal autocorrelation. Surface feeding behaviour could only be analyzed for whale pinger (WP) trials, because very little surface feeding was observed in all phases of the seal scarer (SS) trials.

### Purse-seine trial of the whale pingers

In addition to the individual exposure trials, the whale pingers were also used in a practical application trial on board a capelin purse seine vessel (Börkur NK122) for the 2018 season (January-March) operating out of Neskaupstaður in east Iceland. For the season prior to the trial (January – March 2017), the captain of the vessel kept a log of humpback whale sightings and any encirclements in the net. For the 2018 capelin fishing season, ten pingers were attached to the float line of the purse-seine at a distance of 30-40m from each other, complying with the manufacturer’s recommendations. The captain of the vessel kept record of any issues there were with the use of the pingers, and any incidences of whales inside the net. In addition, one researcher (C.J.B.) joined as an onboard observer for one trip (February 24-28, 2018). During onboard observations, the track of the vessel and whale sightings were recorded in the SpotterPro app (Conserve.IO) during all transit and active fishing days. The number of net casts and tonnes of fish caught with each cast was also noted. Any encirclements of whales with the net were video recorded for documentation using a hand-held video camera (Sony HDR-CX160E handycam).

## Results

### Experimental exposure to acoustic alarms

A total of 23 research trips were undertaken in 2017-2018 totalling approximately 83 hours of effort (Table 1). Of these, enough data for analysis was collected on 14 trips resulting in 9 WP trials and 7 SS trials.

**Table 1.**
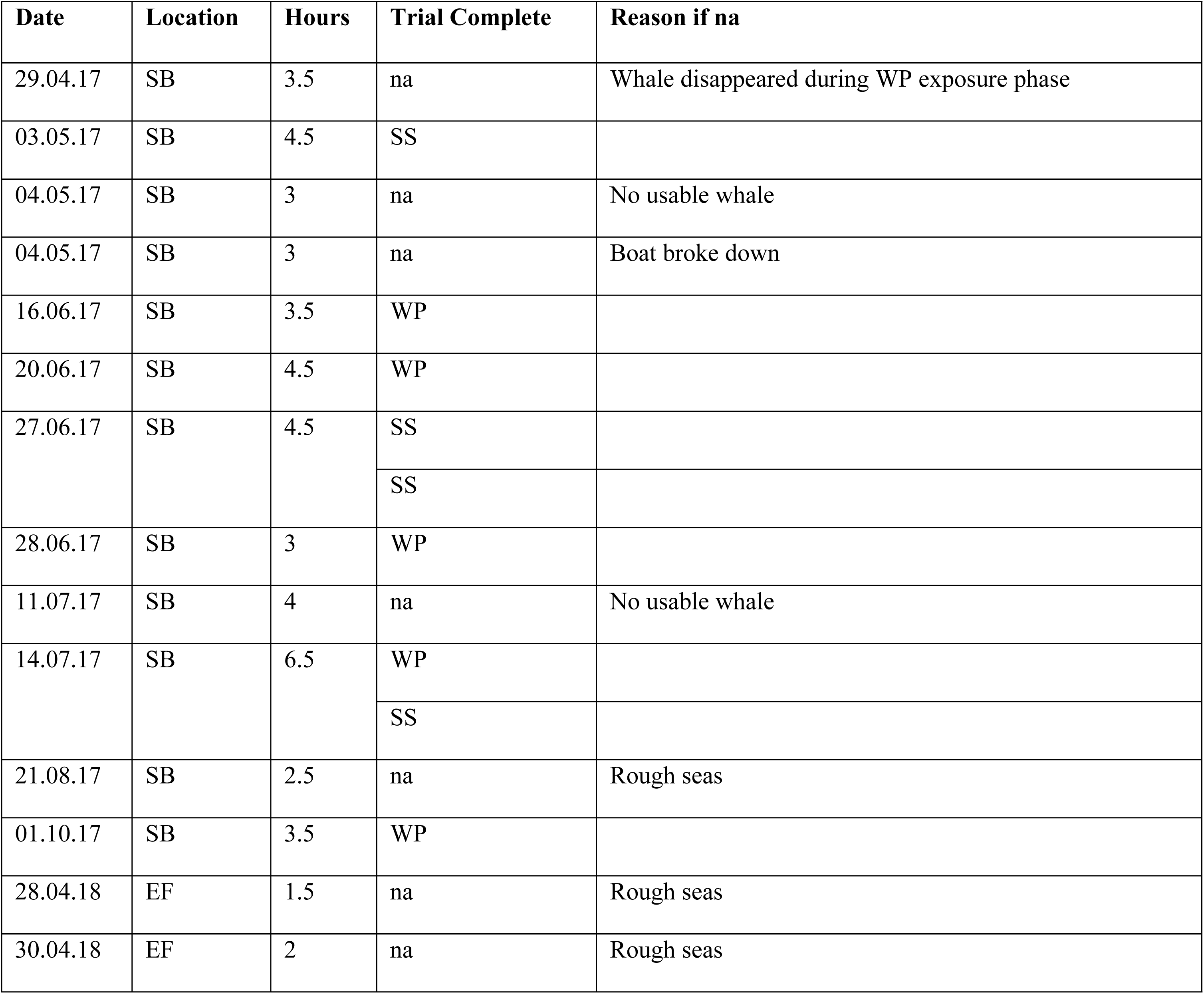
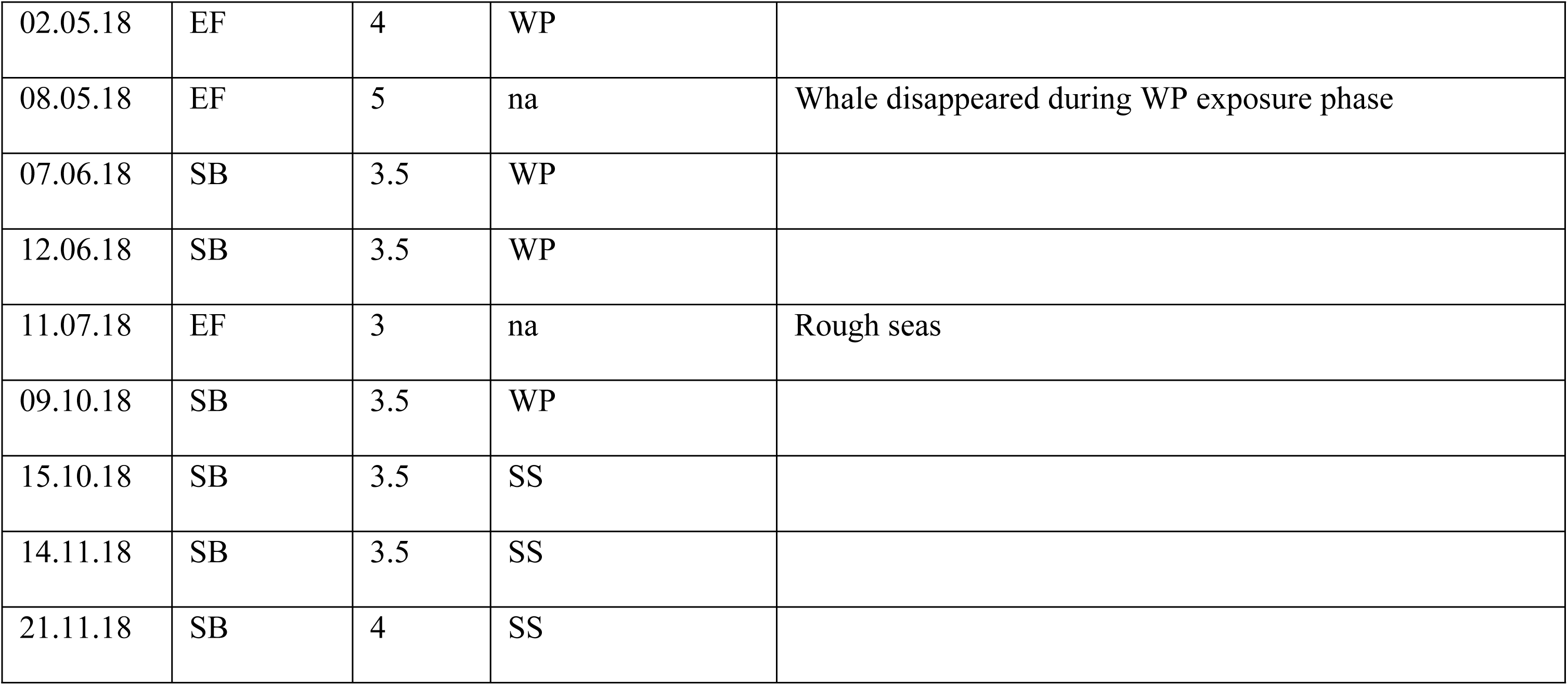
Data collection trips undertaken in 2017-2018 with the Date (DD.MM.YY), Location (SB = Skjálfandi Bay, EF = Eyjafjörður), number of hours (Hours), what trial was completed (Trial Complete: na = not available; no usable trial, SS = seal scarer, WP = whale pinger), and the reason the trip did not result in a usable trial (Reason if na).

| Date | Location | Hours | Trial Complete | Reason if na |
| --- | --- | --- | --- | --- |
| 29.04.17 | SB | 3.5 | na | Whale disappeared during WP exposure phase |
| 03.05.17 | SB | 4.5 | SS |  |
| 04.05.17 | SB | 3 | na | No usable whale |
| 04.05.17 | SB | 3 | na | Boat broke down |
| 16.06.17 | SB | 3.5 | WP |  |
| 20.06.17 | SB | 4.5 | WP |  |
| 27.06.17 | SB | 4.5 | SS |  |
|  |  |  | SS |  |
| 28.06.17 | SB | 3 | WP |  |
| 11.07.17 | SB | 4 | na | No usable whale |
| 14.07.17 | SB | 6.5 | WP |  |
|  |  |  | SS |  |
| 21.08.17 | SB | 2.5 | na | Rough seas |
| 01.10.17 | SB | 3.5 | WP |  |
| 28.04.18 | EF | 1.5 | na | Rough seas |
| 30.04.18 | EF | 2 | na | Rough seas |
| 02.05.18 | EF | 4 | WP |  |
| 08.05.18 | EF | 5 | na | Whale disappeared during WP exposure phase |
| 07.06.18 | SB | 3.5 | WP |  |
| 12.06.18 | SB | 3.5 | WP |  |
| 11.07.18 | EF | 3 | na | Rough seas |
| 09.10.18 | SB | 3.5 | WP |  |
| 15.10.18 | SB | 3.5 | SS |  |
| 14.11.18 | SB | 3.5 | SS |  |
| 21.11.18 | SB | 4 | SS |  |

Fifteen individual whales were used for the successful behavioural trials which produced usable data (Fig 5). Only one individual whale was used twice, in two separate SS trials, but these trials were conducted 18 months apart. Fourteen of the individuals could be identified in the Húsavík Research Center humpback whale catalogues. One individual in Eyjafjorður was not identifiable beyond confirming that it was only used once in the study.

**Fig 5.**
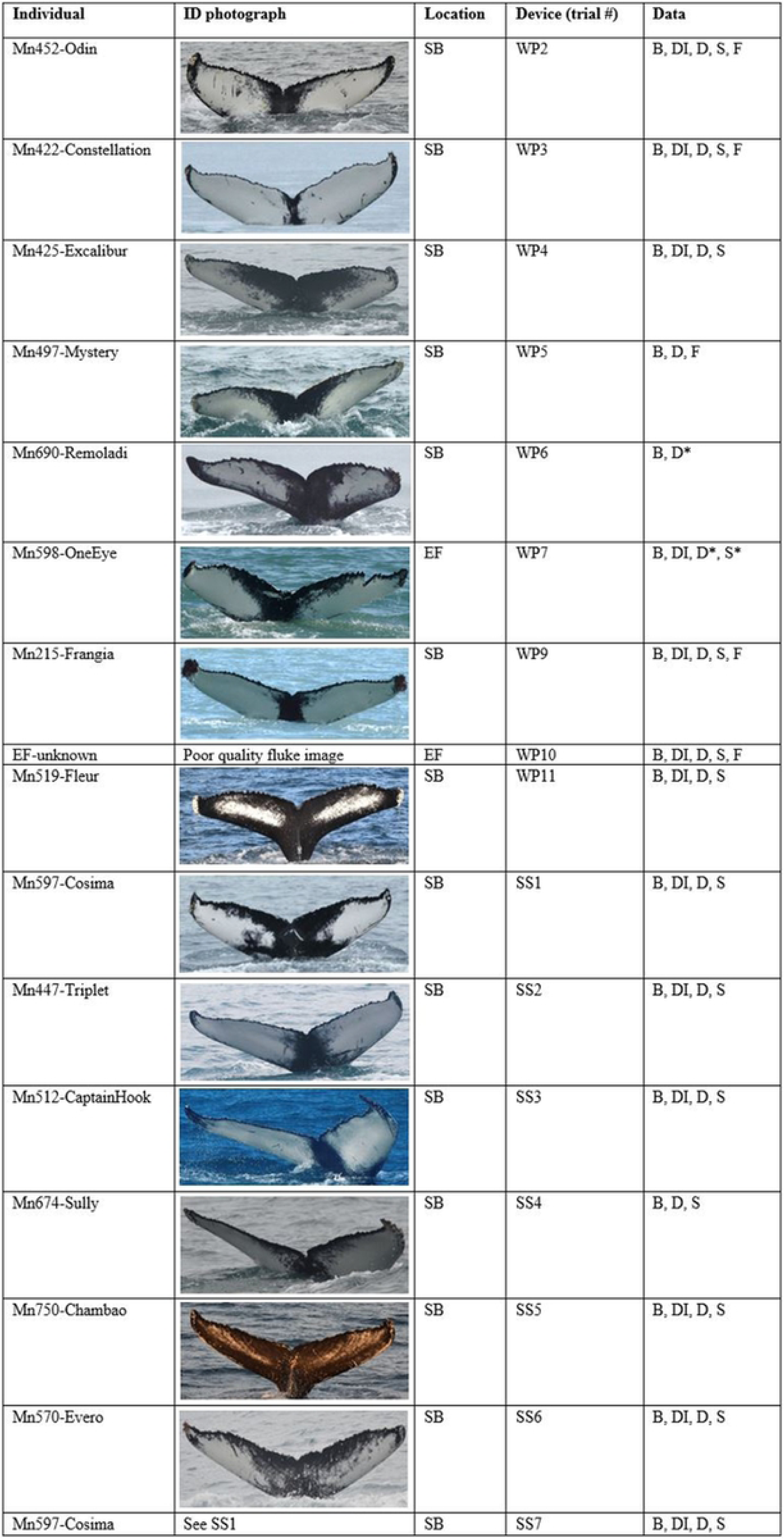
Table showing the individuals used for each successful behavioural trial (expressed by identification code and nickname), location (SB = Skjálfandi Bay, EF = Eyjafjörður), the device used in the trial and the trial ID number (WP = whale pinger, SS = seal scarer), and the data that was collected in each trial (B = breathing rate, DI = directness index, D = dive time, S = swimming speed, F = feeding). Data codes denoted with an * indicate trials for which data is only available for the pre-exposure and exposure phases.

There were eleven attempts made to complete a WP trial, resulting in nine usable trials. Out of these eleven attempts, the individual whale was considered lost (disappeared for more than 20 minutes) in three cases (WP1, WP7, WP8). Two out of these three cases did not result in enough data to be included in the analysis (WP1, WP8). No individuals were lost during SS trials.

Averages of the behavioural response variables for the *PrE*, *E* and *PoE* phases of each WP trial are shown in Fig 6. Full models for each behavioural response variable included the experimental phase (*PrE*, *E*, *PoE*) as fixed effect and a random intercept and slope (for experimental phase) in addition to an autoregressive correlation structure of order 1 as random effects. Random slope models did not fit the data significantly better than random intercept models for the behavioural response variables speed, surface feeding, breathing rate and directness (Table 2). Thus, there was no statistical support for individual variation in these responses.

**Fig 6.**
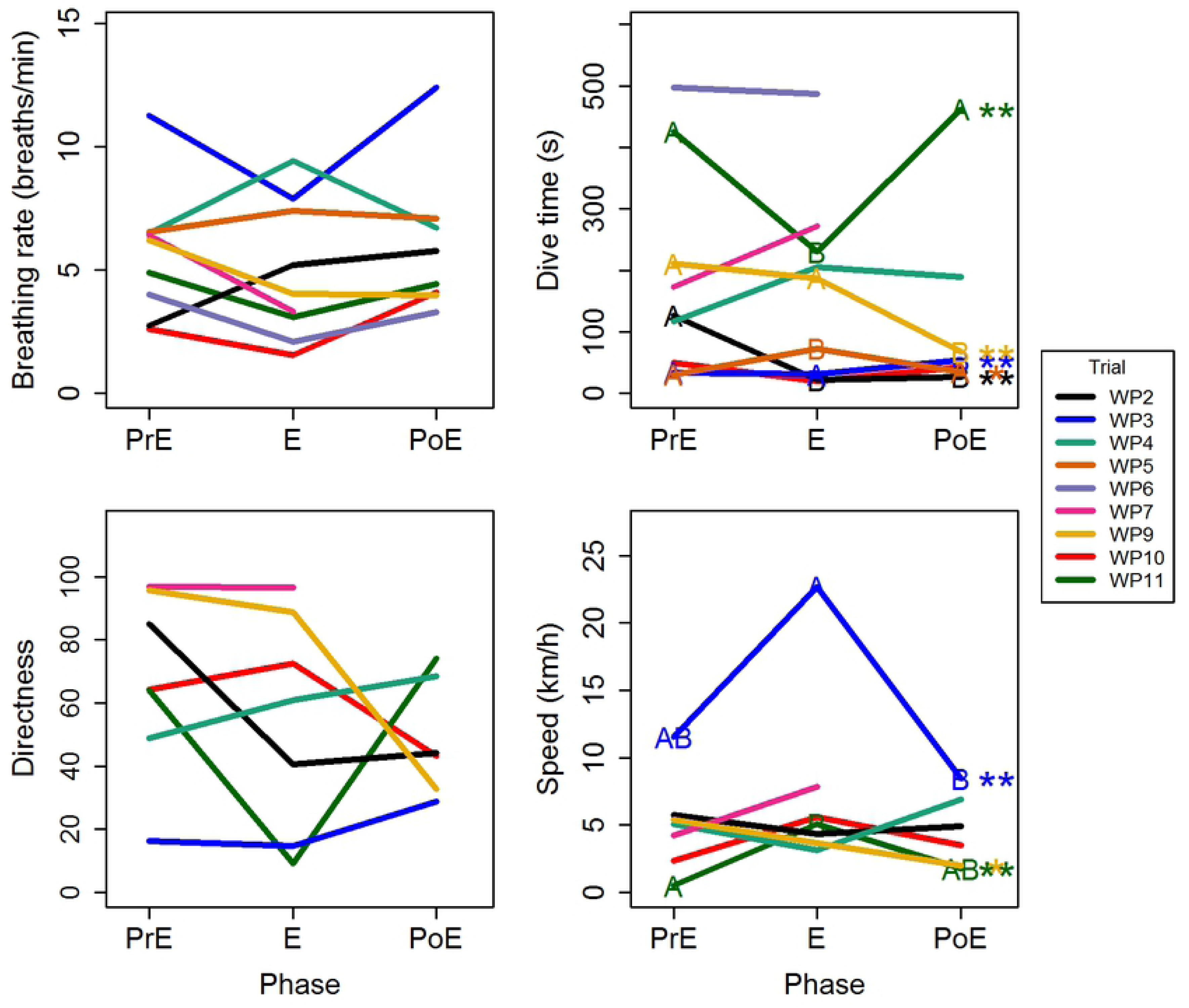
Averages of the behavioural response variables breathing rate, dive time, directness and speed for the pre-exposure (PrE), exposure (E) and post-exposure (PoE) phases of each whale pinger (WP) trial. Stars highlight individual whale pinger trials in which the response variable differed significantly between the phases (* uncorrected p < 0.05; ** Bonferroni-corrected p < 0.05). Letters indicate between which phases significant differences occurred. Models for individual whale pinger trials were only calculated for response variables for which overall models found a significant effect of phase or random slope (see Table 3).

**Table 2.**
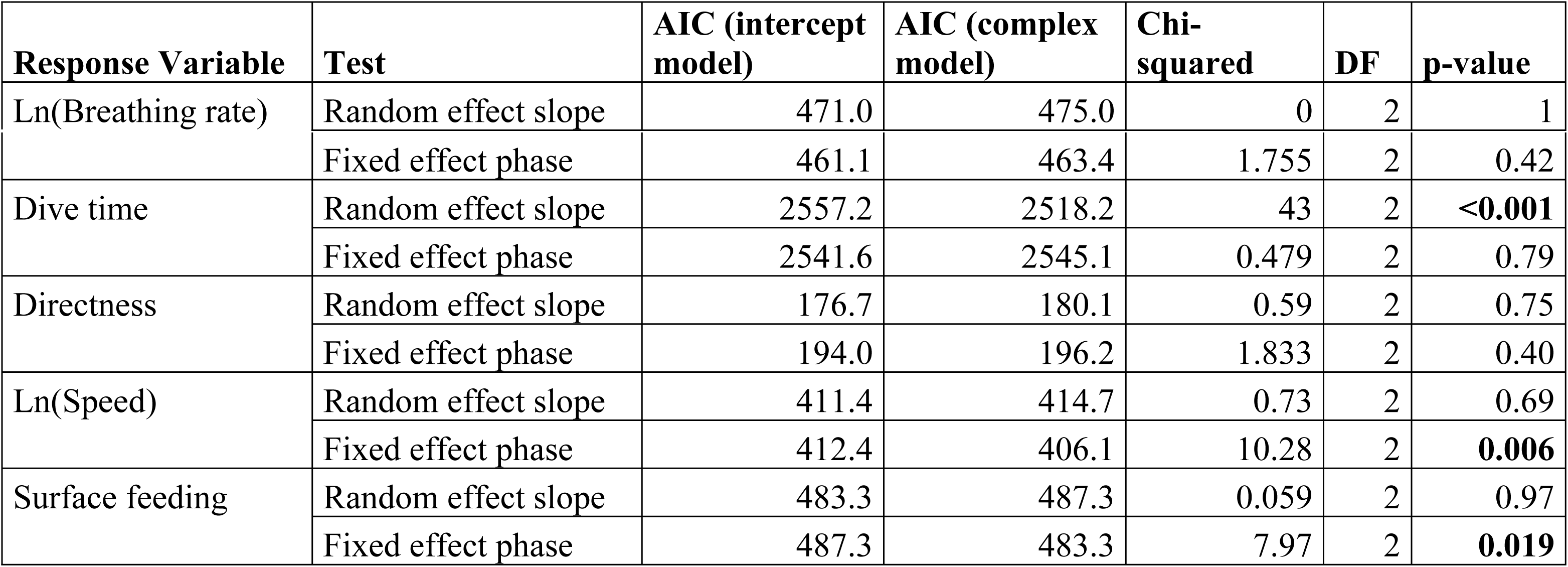
Assessment of the random and fixed effects structures of five models explaining the change in a behavioural response variable after exposure to a whale pinger. To test if the effect sizes of the contrasts to the pre-exposure phase differed significantly between individuals, a random intercept and slope model was compared to a pure random intercept model by means of comparison of Akaike Information Criterion (AIC) values and a likelihood ratio test. The fixed effects structure was tested by comparing models with and without the predictor phase. Assessment of random effects was based on models estimated by restricted maximum likelihood, whereas assessment of fixed effects was based on maximum likelihood estimation. Significant p-values are bolded.

| Response Variable | Test | AIC (intercept model) | AIC (complex model) | Chi-squared | DF | p-value |
| --- | --- | --- | --- | --- | --- | --- |
| Ln(Breathing rate) | Random effect slope | 471.0 | 475.0 | 0 | 2 | 1 |
|  | Fixed effect phase | 461.1 | 463.4 | 1.755 | 2 | 0.42 |
| Dive time | Random effect slope | 2557.2 | 2518.2 | 43 | 2 | **<0.001** |
|  | Fixed effect phase | 2541.6 | 2545.1 | 0.479 | 2 | 0.79 |
| Directness | Random effect slope | 176.7 | 180.1 | 0.59 | 2 | 0.75 |
|  | Fixed effect phase | 194.0 | 196.2 | 1.833 | 2 | 0.40 |
| Ln(Speed) | Random effect slope | 411.4 | 414.7 | 0.73 | 2 | 0.69 |
|  | Fixed effect phase | 412.4 | 406.1 | 10.28 | 2 | **0.006** |
| Surface feeding | Random effect slope | 483.3 | 487.3 | 0.059 | 2 | 0.97 |
|  | Fixed effect phase | 487.3 | 483.3 | 7.97 | 2 | **0.019** |

**Table 3.**
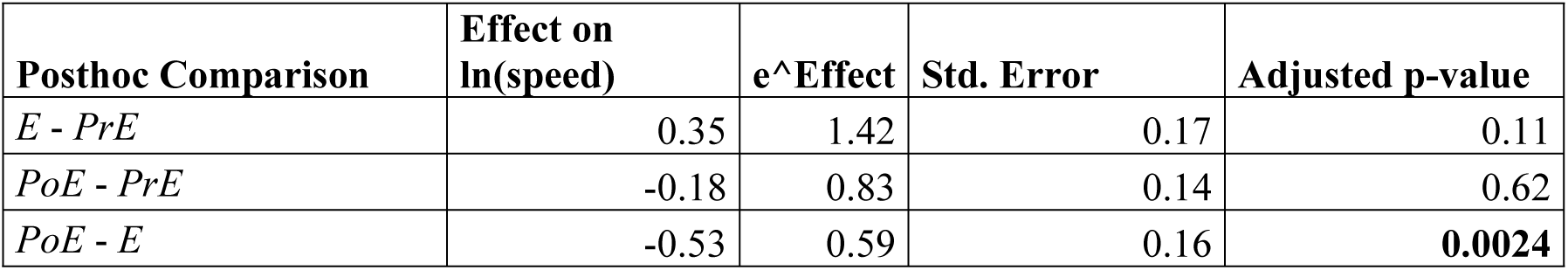
Posthoc comparison for the predictor phase (*PrE* = pre-exposure, *E* = exposure, *PoE* = post-exposure) in the swimming speed model based on the whale pinger data (See Table 2). Since the response variable speed is ln-transformed, effect is the difference in ln(speed) and e^Effect is the ratio between speeds in the two compared phases. Adjusted p-values are Bonferroni-corrected p-values. Significant p-values are bolded.

The predictor phase had a significant effect on both speed (p = 0.006; Table 2) and surface feeding (p = 0.019; Table 2). Humpback whale speed during the *E* phase was 1.7 times higher than during the *PoE* phase (p = 0.0024; Table 3) and 1.4 times higher than during the *PrE* phase (p = 0.11; Table 3). No significant differences in humpback whale speed were observed between the *PrE* and *PoE* phases (p = 0.62; Table 3). The probability of surface feeding was significantly lower during the *E* phase than during the PoE phase (p = 0.026; Table 4). The reduction in surface feeding from the *PrE* to the *E* phase was marginally significant (p = 0.099; Table 4). Rates of surface feeding amounted to 11% and 13% in the *PrE* and *PoE* phases and dropped to 4% in the *E* phase (Fig 7).

**Fig 7.**
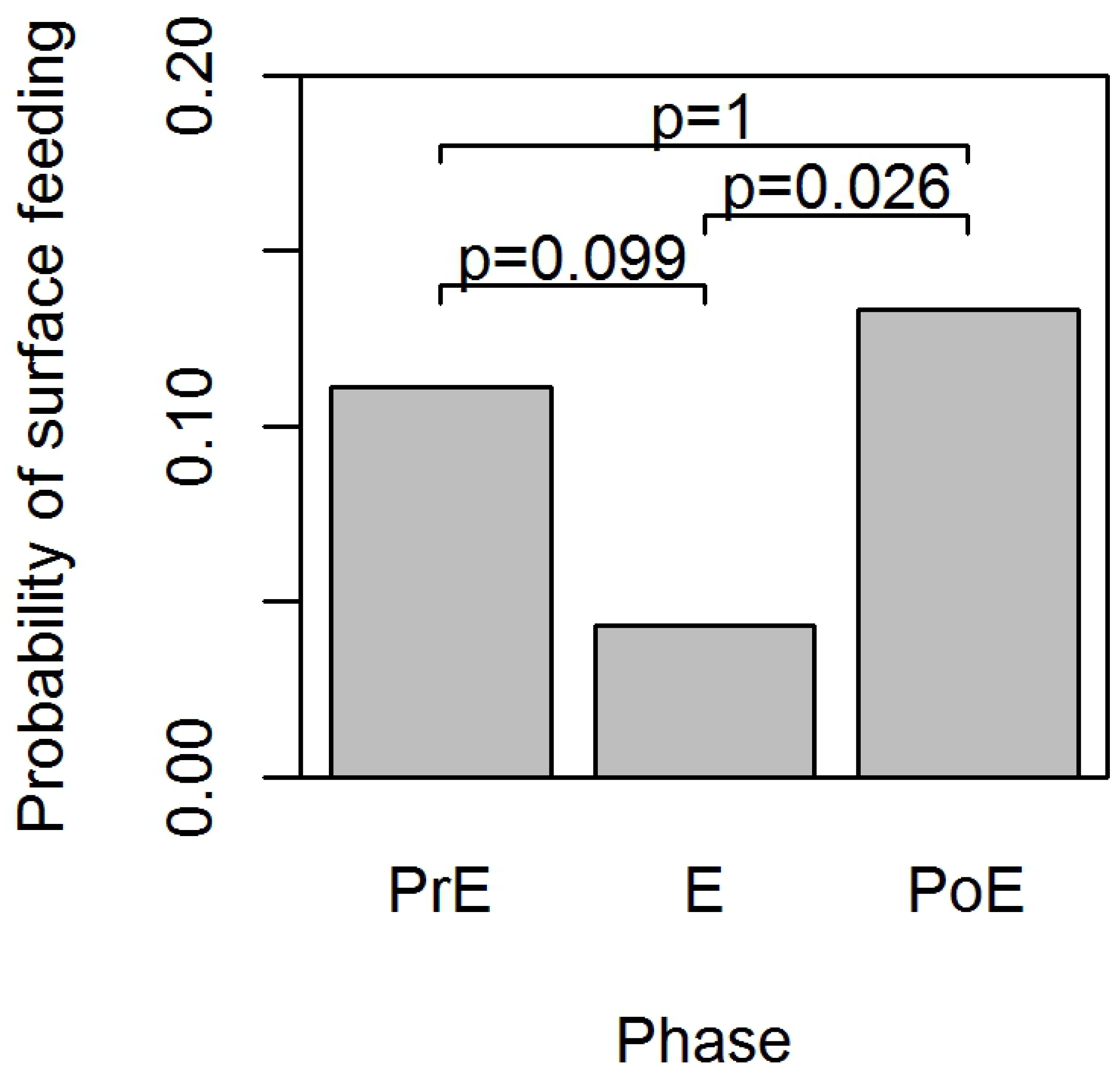
Graph showing the probability of surface feeding during whale pinger trials for each phase (*PrE* = pre-exposure phase, *E* = exposure phase, *PoE* = post-exposure phase). P-values are Bonferroni-corrected p-values obtained in the posthoc comparison (See Table 4).

**Table 4.** Posthoc comparison for the predictor phase (*PrE* = pre-exposure, *E* = exposure, *PoE* = post-exposure) in the surface feeding model based on the whale pinger data (See Table 2). Effect and std. error are the effect size on the linear predictor scale and its standard error. Adjusted p-values are Bonferroni-corrected p-values. Significant p-values are bolded.

| Posthoc Comparison | Effect on surface feeding | Std. Error | Adjusted p-value |
| --- | --- | --- | --- |
| *E* - *PrE* | -1.02 | 0.48 | 0.099 |
| *PoE* - *PrE* | 0.21 | 0.26 | 1 |
| *PoE* - *E* | 1.22 | 0.47 | **0.026** |

No significant changes in breathing rate (p = 0.42; Table 2) and directness (p = 0.40; Table 2) were detected in response to exposure to whale pinger sound. The model for dive time was the only case in which a random slope model fitted the data significantly better than a random intercept model (p < 0.001; Table 2). Phase of the trial, however, had no significant effect on dive time (p = 0.79; Table 2).

The averages of the behavioural response variables for the *PrE*, *E* and *PoE* phase of each SS trial are displayed in Fig 8. Models of the behavioural response variables were set up in the exact same manner as for the WP trial analysis. Random slope models did not fit the data significantly better than random intercept models for any of the response variables (Table 5). Experimental phase did not have a significant effect on any of the response variables (Table 5). Thus, we found no evidence for an individual-specific or shared response of humpback whales to seal scarer alarm.

**Fig 8.**
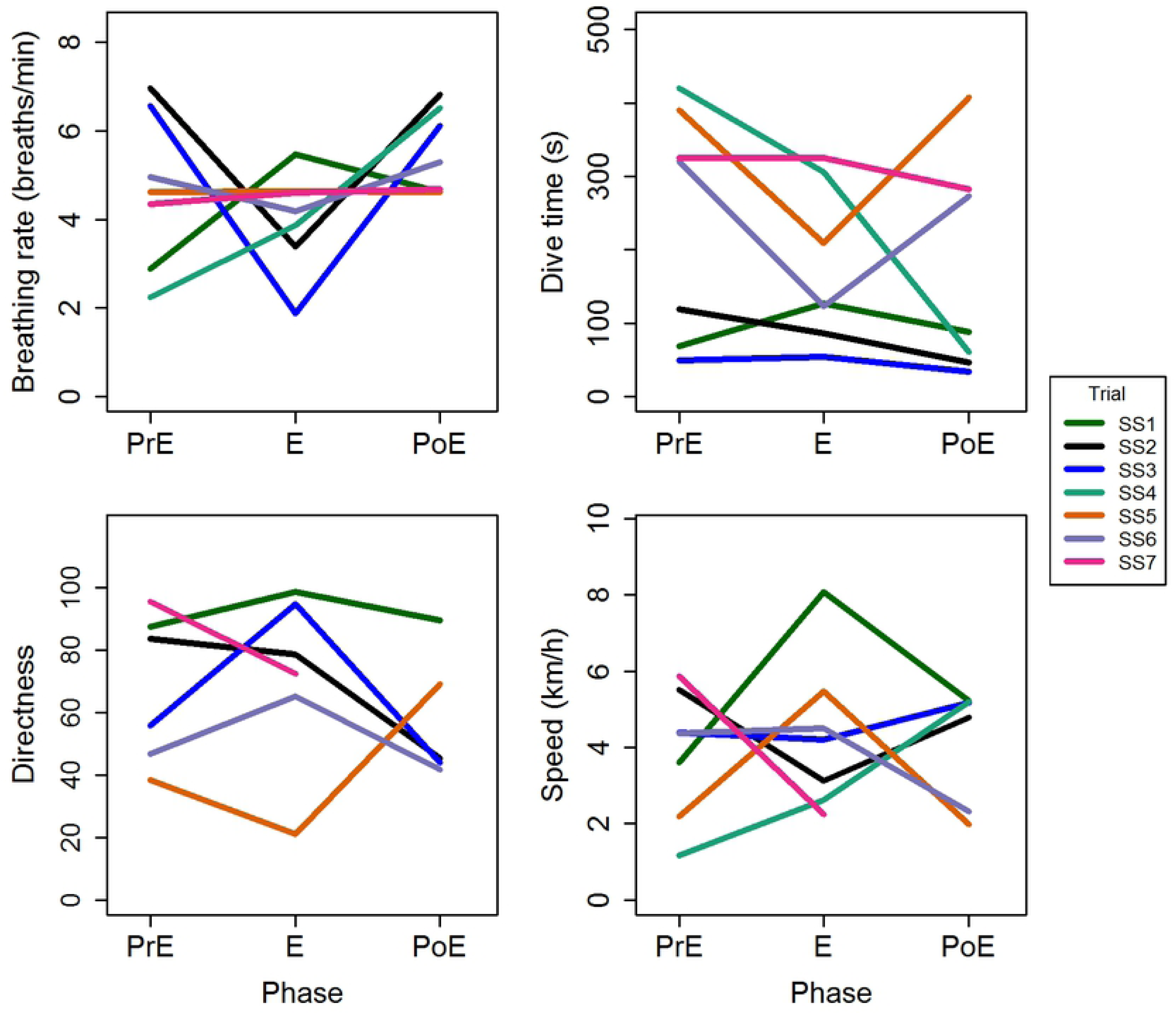
Averages of the behavioural response variables breathing rate, dive time, directness and speed for the pre-exposure (PrE), exposure (E) and post-exposure (PoE) phases of each seal scarer (SS) trial.

**Table 5.**
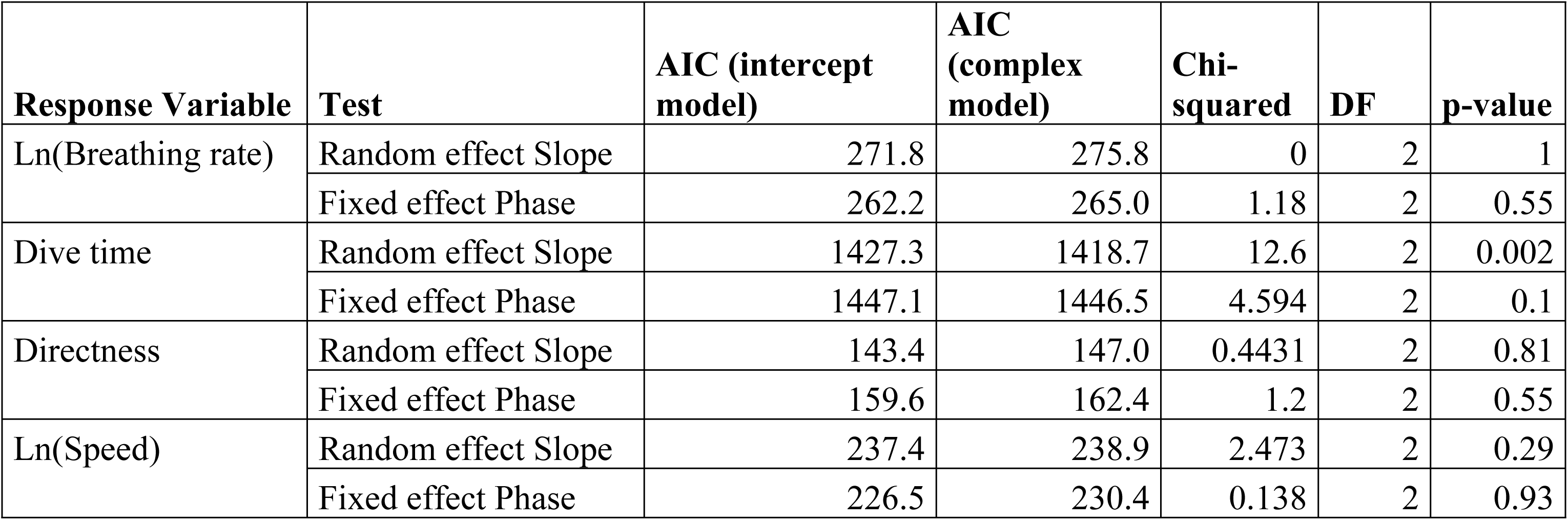
Assessment of the random and fixed effects structures of five models explaining the change in a behavioural response variable after exposure to a seal scarer. To test if the effect sizes of the contrasts to the pre-exposure phase differed significantly between individuals, a random intercept and slope model was compared to a pure random intercept model by means of comparison of Akaike Information Criterion (AIC) values and a likelihood ratio test. The fixed effects structure was tested by comparing models with and without the predictor phase. Assessment of random effects was based on models estimated by restricted maximum likelihood, whereas assessment of fixed effects was based on maximum likelihood estimation.

### Purse-seine trial of the whale pingers

The captain of the participating capelin purse seine vessel did not report any issues with humpback whales inside the net in the 2017 season and reported that there were generally lower sightings and incidences than in the previous (2016) season. During the 2018 capelin fishing season, the onboard observer recorded 34 individual humpback whale sightings at 7 locations during 16 hours of observation (Table 6) with 70.6% (n = 24) occurring while the boat was in the capelin fishing grounds off the south/southwest coast of Iceland. The net was cast 3 times during onboard observation and a total of 1510 tonnes of capelin were caught.

**Table 6.** Effort during onboard observation on the capelin purse seine vessel using the whale pingers including date (DD,MM,YY), time, whale sightings (Mn = humpback whale (*Megaptera novaeangliae*), Bp = fin whale (*Balaenoptera physalus*), location (latitude, longitude), status of the boat, and comments. A * denotes in which sighting two whales were encircled in the net.

| Date | Time | Sightings | Boat Location | Status | Comments |
| --- | --- | --- | --- | --- | --- |
| 24.02.18 | 20:00 |  |  | leaving port |  |
| 25.02.18 | 9:00-12:00 | 1Mn | 63.514593N -<br>17.864615W | transit |  |
| 25.02.18 | 13:40-15:40 | NA |  | transit |  |
| 25.02.18 | 16:10-18:10 | 4Mn | 63.430734N, -<br>19.595009W | transit/docking | Two pairs of whales |
|  |  |  | 63.437207N, -<br>19.901842W |  |  |
| 26.02.18 |  |  |  | in port |  |
| 27.02.18 | 9:30-11:45 | 9Mn | 63.722643N, -<br>20.818695W | transversing grounds |  |
|  |  |  | 63.727772N, -<br>20.836443W |  |  |
|  |  |  | 63.736444N, -<br>20.884974W |  |  |
|  |  |  | 63.737366N, -<br>20.887947W |  | Pair of whales |
|  |  |  | 63.75826N, -<br>20.912274W |  |  |
|  |  |  | 63.764711N, -<br>20.92963W |  | Pair of whales |
|  |  |  | 63.785706N, -<br>20.989416W |  |  |
| 27.02.18 | 13:30-16:00 | 3Mn | 63.766879N, -<br>20.969197W | transversing<br>grounds/fishing |  |
|  |  |  | 63.729173N, -<br>20.892273W |  |  |
|  |  |  | 63.784402N, -<br>20.985329W |  |  |
| 27.02.18 | 17:00-18:00 | 3Mn | 63.604162N, -<br>20.773594W | transversing grounds | Pair of whales |
|  |  |  | 63.596906N, -<br>20.714554W |  |  |
| 28.02.18 | 8:35-10:00 | 6Mn* | 63.499573N, -<br>20.940331W | fishing | Pair of whales |
|  |  |  | 63.498824N, -<br>20.937591W |  |  |
|  |  |  | 63.498249N, -<br>20.945592W |  |  |
|  |  |  | 63.500046N, -<br>20.946389W |  |  |
|  |  |  | 63.499475N, -<br>20.9425W |  |  |
| 20.02.18 | 16:45-18:00 | 5Mn | 63.372122N, -<br>18.879468W | transit |  |
|  |  |  | 63.36858N, -<br>18.812109W |  |  |
|  |  |  | 63.367307N, -<br>18.774776W |  |  |
|  |  |  | 63.378524N, -<br>18.599105W |  |  |
|  |  |  | 63.382446N, -<br>18.559984W |  |  |
|  |  | 1Bp | 63.369304N, -<br>18.693336W |  |  |

Whales at the surface near the vessel when fishing operations were beginning were noted to swim away from the area, with one whale specifically observed turning 180 degrees to go opposite of where the seine net was being set into the water. There were two incidences where humpback whales were encircled in the net fitted with the whale pingers during 2018, once when the onboard observer was present and once when they were not. In both incidences two humpback whales appeared at the surface inside the net once the bottom of the net was being closed, indicating the whales entered from the bottom. During the onboard observation incident, it was noted the whales were “trumpeting” and showed signs of being distressed by the encirclement in the net. In an attempt to release the whales in both cases, the extra line attaching the end of the net to the vessel, without floats or pingers attached, was not brought in towards the boat while the net was closed at the bottom creating an approximately 100m wide opening in the side of the net towards the stern. During the onboard observation encirclement, the two whales spent approximately 5 minutes inside the net before locating the opening and escaping without causing any damage. According to the captain the second incident occurred in the exact same manner. The captain and crew reported that whales rarely, if ever, find this opening and escape without further action or damage to the net in previous seasons when the pingers were not in use. Only 270 tonnes of capelin were caught in the cast where the whales were encircled in the net during onboard observation (compared to 690 and 550 tonnes in the other two casts).

## Discussion

Mitigating large whale entanglement in fishing industries is of global interest. This study represents the first in situ experiments testing commercially available acoustic alarms on humpback whales in their North Atlantic feeding grounds off Iceland, and the first study to consider feeding as a behavioural response variable. Results showed that it was significantly less likely to observe surface feeding behaviour during the *E* phase of the WP trials, when the whale pinger was active in the water, compared to when the whales were observed prior to and after exposure. This suggests that the whales reduce or stop surface feeding in response to the pinger. Previous studies have found that humpback whales cease feeding in response to sonar sounds [59], decrease side roll feeding in response to ship noise [60], and decrease detectable lunge feeding behaviour during approaches of whale watching vessels in one of the study sites for this experiment (Skjálfandi Bay) [61], suggesting reduction in feeding may be a common response when whales are exposed to anthropogenic noise. This is the first time a reduction in feeding behaviour has been documented in response to a pinger alarm and we can only hypothesize why the whales would react this way. One possibility is that they are simply distracted by or curious about the sudden introduction of an unnatural and unfamiliar sound in their environment. Since the received sound level from the whale pinger was likely low, it is unlikely that the whales were startled and stopped feeding, and there was no clear indication that they moved away from the sound based on results from the directness index model. However, three out of the eleven individuals involved in attempted WP trials were declared lost (disappeared for more than 20 minutes) during the *E* phase when the whale pinger was in the water. Two out of the three cases (WP1, WP8) did not have enough data to include them in analysis. The first individual (WP1) took one dive approximately 200m away after the *E* phase started and then never resurfaced in sight. The second individual (WP7), which was included in analysis, started traveling and stopped diving with the fluke in the air shortly after the *E* phase began and was last sighted an estimated 1000m away before it was lost. The third individual (WP8) was last seen before the pinger was put in the water to begin the *E* phase and was not seen again within 20 minutes after the *E* phase began. Since the boat was stationary during the *E* phase of the trials, the probability of losing sight of the whale is higher than during the *PrE* and *PoE* phases when the boat is maneuvered. However, trials were only conducted in good weather with good visibility and therefore the complete disappearance in the 15-minute *E* phase was most likely due to a change in behaviour. It is possible that these individuals were disturbed by the pinger sound and moved away.

Lien et al. [27] reported that there was an increase in cod catch in traps that had alarms attached than those that did not, suggesting target fish species are not affected by the alarms. Humpback whales are primarily feeding on smaller fish species in the North Atlantic, such as capelin [62]. Fish are modelled to hear at low frequencies below 0.5-1 kHz and react to high intensity sound [63], therefore it is unlikely that the pinger sound affected the prey that the whales were feeding on during the trials. This suggests that the whales responded to the pinger sound directly rather than to a change in prey distribution or behaviour.

Whales reducing or stopping their feeding in response to the pinger, could lead to lower incidence of humpback whale encirclement and entanglement in Icelandic fisheries since the whales are likely feeding when these incidences occur. Humpback whales in coastal polar waters have been recorded making an average of 28 feeding lunges per hour in Antarctica, and a tagged whale in the primary field site for this study (Skjálfandi Bay) was similary recorded making an average of 33 feeding lunges per hour [61], suggesting the majority of their time is spent foraging. Furthermore, entanglement of humpback whales has been observed as coinciding with the spawning of one of their main prey species, capelin, in Newfoundland Canada [65] and encirclement of humpbacks in Iceland that were evidently feeding on capelin at the time of the incident was observed during this study. We therefore hypothesize that if the whales stop feeding in the vicinity of fishing gear with active pingers they may be more likely to take notice of the gear and less likely to become entangled or encircled. Therefore, the pingers may be a useful mitigation tool. The whales that were encircled in the purse seine net using the pingers in this study were not surface feeding and entered the net from deeper than 120m while the pingers were near the surface of the water, which may indicate the pingers were not in the correct position to cause the whales to stop feeding and avoid entering the net. Overall, this suggests that if the whales stop feeding in response to the pinger, it may reduce the risk of them becoming entangled or encircled in the fishing gear, but the pingers need to be positioned strategically on the net at the appropriate depth to elicit the reduced feeding response. Further experimentation with the pingers at different depths and tagging of the whales in order to have information about their underwater feeding activity could provide valuable information for this hypothesis.

Disruption of feeding behaviour in these whales is cause for concern for negative impacts on the individual, and possibly the population, if pinger use becomes widespread in the fishing industry. Humpback whales need to consume an estimated 1432 Kcal of food per day during the summer feeding season in order to have a large energy storage for their migration and winter breeding season [39]. Insufficient energy stores may lead to decreased ability to migrate or decreased reproductive success, which can furthermore impact the recruitment rate of the population [66]. However, it is important to note that exposure to the whale pinger during the *E* phase was only for 15 minutes, so it is unknown if the whales would habituate to the sound and continue feeding normally after a longer period of time. We did not observe any lasting effect of the whale pinger on surface feeding, suggesting that when the pinger is removed from the water the whale quickly returns to its post-exposure behaviour. Further investigation into the humpback whale’s feeding response to low frequency acoustic alarms is recommended for the future in order to determine if this response is consistent within larger sample sizes and if it is detected in other humpback whale feeding grounds. It is also advisable to investigate what the response of the whales is to longer exposure to the alarms to determine if reduction in feeding is only a short-term consequence or is a longer response.

Given the uncertainty of the effects of long-term use, the whale pingers may be particularly advisable for fishing methods in which the gear is not in the water for long periods of time such as attached to purse seine nets or suspended in the water from long-line vessels.

The whales also significantly increased their swimming speed during exposure to the whale pinger. An increase in humpback whale swimming speed has been documented in response to whale watching boats [67, 68], but has not been reported in previous studies investigating behavioural responses to pingers. Boye et al. [67] found that whales took significantly shorter dives and increased their mean speed in response to boats, while similarly in our study some individual whales significantly decreased their dive time, while overall the whales significantly increased their speed. The increase in speed supports that humpback whales respond to the whale pinger sound. However, further investigation into the whales’ behaviour underwater is needed to infer the effect of this reaction on entanglement mitigation.

Though similar previous studies were conducted during whale migration opposed to during the feeding season, results from this study were consistent with recent previous finding that there is no consistent, significant behavioural response of humpback whales to the whale pinger in terms of dive time, breathing rate, or directness [24, 25, 26]. There was also no evidence for individual-specific responses in terms of breathing rate or directness, meaning we have no evidence that individuals reacted to the pinger significantly in terms of these variables. The received sound level may have been too low to elicit a detectable behavioural change in terms of these variables, which are behavioural reactions that can indicate the whale was disturbed or startled [28]. The humpback whales foraging in the study sites for this study are also regularly exposed to a lot of anthropogenic noise. Both locations host a high number of whale-watching vessels which target humpback whales primarily for their sightings, as well as industrial ports with associated development and maintenance noise and fishing vessels, cruise ships and cargo ships entering and exiting often. There are also commercial fishing grounds within the waters of both study sites. This may mean that many humpback whales in this area are generally habituated to anthropogenic noise and may not show behavioural changes that would indicate they are significantly disturbed or stressed.

There was evidence for individual-specific responses in terms of dive time, even though there was no significant effect of whale pinger exposure on dive time overall. Some individuals significantly increased dive time during the *E* phase while other individuals significantly decreased. This could indicate that individuals just had variable significant reactions when the sound was introduced, which could depend on their behavioural state in the *PrE* phase, as suggested by Southall et al. [54]. It is also possible that individuals were just naturally changing between behavioural states from a long dive period to a short dive period and vice versa during the trials and for some individuals this happened to coincide with the *E* phase of the trial. Further investigation into humpback whale dive time response to low frequency sound, taking into account initial behavioural state and natural changes in behaviour, are necessary to conclude whether dive time changes are in response to the pinger or not.

We found no evidence for a significant effect of the seal scarer alarm on humpback whale speed, dive time, breathing rate or directness. In addition, we found no evidence that there were any individual-specific responses to the seal scarer in terms of any of these variables. The seal scarer was measured as having a source level 52 dB louder than the whale pinger and due to this it was hypothesized the whales would have some reaction to the loud sound even though the frequency of the alarm is at the top or slightly above the estimated hearing range of the humpback whales [40]. Hearing in minke whales has more recently been modelled using CT scanning to show their range is higher than what is necessary for their communication [69], and they showed significant behavioural reactions to the seal scarer in Iceland [33], but we found no evidence that this is similar for the humpback whales. This is consistent with the findings of Henderson [70] who also concluded humpback whales do not react to high frequency pingers, though the pinger used in their study was 17-45 kHz higher in frequency than the device used in our study. It is possible that the frequency of the seal scarer was just too high for the humpback whales to hear the alarm well enough to exhibit a significant response, confirming that acoustic entanglement mitigation devices need to target the best-estimated hearing range of the whales. However, the surface feeding behavioural response remains unknown for the seal scarer since there was not enough surface feeding observed in the trials to analyze this.

The use of the whale pingers on the capelin purse seine net for one season provided a first insight into the use of the devices in a practical application in Iceland. A pair of whales entered the net fitted with the pingers from the bottom, before it had been closed, twice. Since the pingers were attached along the float line at the top of the net, this made sense that whales may still enter from the bottom, with the net extending down approximately 120m. Despite this, in both cases the whales were able to find their way out of the net through an approximately 100m wide (at the surface) opening to the side of the net without causing any damage and without further intervention methods from the captain (such as putting the boat into reverse to sink the float line), a very rare occurrence according to the captain and crew onboard. This led to an overall positive view of the whale pingers and an increased interest in further trials for use in the Icelandic capelin purse seine fishery to prevent net damage.

Suggestions for repositioning the pingers on the net could be considered in the future including attaching the pingers to the lead line at the bottom of the net or sewing specialized pockets for the pingers into the lower portion of the net (Hjörvar Hjálmarsson pers. comm. 2018, Geir F. Zoega pers. comm. 2019). These observations also led to hypothesizing about the currently unknown directional hearing capabilities of humpback whales. Ten pingers were spaced approximately every 30-40m along the net measuring 450×120m in total. When the whales were inside the net there was an approximately 100m opening left at the surface by a single rope attaching the net end to the vessel, and the first pinger was attached to the net approximately 30m from the “bag” netting (the net that remains in the water to prevent fish from escaping as the they are hauled on board). This equals an estimated 150m pinger-less space, of which approximately 100m is the opening for the whales to escape through. If the pingers were truly aiding in the humpback whales finding this opening, as is being suggested based on the captain and crew’s experience with whales becoming encircled in the net for several years, this suggests that the whales were able to acoustically detect this 150m pinger-less space where the sound level was lower, and then find the 100m opening. Further trials and observation of whale pinger use on purse seine nets could provide more insight into this hypothesis.

Low frequency whale pingers may be a useful tool in preventing humpback whale entanglement and net damage occurring in their feeding grounds based primarily on the findings from this study that the whales reduced their surface feeding behaviour in response to the pinger and exited a purse seine net equipped with pingers without net damage or intervention. The whale pinger also had a significant effect on the swimming speed of the whales in this study, however the implications of this response in terms of entanglement reduction are unknown. The fact that we observed no consistent behavioural reaction to the whale pinger in terms of dive time, breathing rate or directness suggests that the whale pingers do not elicit a stress response in humpback whales although whales increase swimming speed and reduce feeding. No significant reactions to the louder, high-frequency seal scarer alarm in terms of speed, dive time, breathing rate or directness were observed which suggests these alarms are not effective for humpback whales, though their feeding response to such an alarm requires further investigation. Though the whale pingers may be effective in mitigating entanglement, they should be used with caution until further information is known about the longer-term consequences of the reduction in feeding, and may be best suited only for certain, short-term applications in conjunction with other possible entanglement mitigation methods such as seasonal or area restrictions on fishing, and modified fishing gear.

## Acknowledgments

A special thank you to Gentle Giants Whale Watching, the boat captains and the internship students at the Húsavík Research Centre who assisted with data collection. In addition, we greatly thank the captain and crew of Börkur for their participation in this study.

## References

1. Moore MJ, Van Der Hoop J, Barco SG, Costidis AM, Gulland FM, Jepson PD, et al. Criteria and case definitions for serious injury and death of pinnipeds and cetaceans caused by anthropogenic trauma. Dis Aquat Organ. 2013; 103(3): 229–264. https://doi.org/10.3354/dao02566

2. Cassoff RM, Moore KM, McLellan WA, Barco SG, Rotstein DS, Moore MJ. Lethal entanglement in baleen whales. Dis Aquat Organ. 2011; 96(3): 175–185. https://doi.org/10.3354/dao02385

3. Knowlton AR, Kraus SD. Mortality and serious injury of northern right whales (*Eubalaena glacialis*) in the western North Atlantic Ocean. J Cetacean Res Manag. (Special Issue). 2001; 2: 193–208.

4. van der Hoop J, Corkeron P, Moore M. Entanglement is a costly life-history stage in large whales. Ecol Evol. 2017; 7(1): 92–106. https://doi.org/10.1002/ece3.2615

5. Moore MJ, van der Hoop JM. The Painful Side of Trap and Fixed Net Fisheries: Chronic Entanglement of Large Whales. J Mar Biol. 2012; 1–4. https://doi.org/10.1155/2012/230653

6. Robbins J, Knowlton AR, Landry S. Apparent survival of North Atlantic right whales after entanglement in fishing gear. Biol Conserv. 2015; 191: 421–427. https://doi.org/10.1016/j.biocon.2015.07.023

7. Volgenau L, Kraus SD, Lien J. The impact of entanglements on two substocks of the western North Atlantic humpback whale, *Megaptera novaeangliae*. Can J Zool. 1995; 73(9): 1689–1698. https://doi.org/10.1139/z95-201

8. Johnson A, Salvador G, Kenney J, Robbins J, Kraus S, Landry SCP, et al. Fishing Gear Involved in Entanglements of Right and Humpback Whales. Mar Mamm Sci. 2005; 21(4): 635–645

9. Gall SC, Thompson RC. The impact of debris on marine life. Mar Pollut Bull. 2015; 92(1–2): 170–179. https://doi.org/10.1016/j.marpolbul.2014.12.041

10. Lien J, Aldrich D. Damage to the inshore fishing gear in Newfoundland and Labrador by whales and sharks during 1981. CAFSAC Marine Mammal Committee Meetings, St. John’s NFL, 1982 May 18-19.

11. Lien J. A study of entrapment in fishing gear: Causes and Prevention. Progress Report 1979 March 1.

12. Erbe C, McPherson C. Acoustic characterisation of bycatch mitigation pingers on shark control nets in Queensland, Australia. Endanger Species Res. 2012; 19(2): 109– 121. https://doi.org/10.3354/esr00467

13. Kraus SD, Read AJ, Solow A, Baldwin K, Spradlin T, Anderson E, et al. Acoustic alarms reduce porpoise mortality. Nature. 1997; 388: 525

14. Jefferson TA, Curry BE. Acoustic methods of reducing or eliminating marine mammal-fishery interactions: Do they work? Ocean Coast Manag. 1996; 31(1): 41– 70. https://doi.org/10.1016/0964-5691(95)00049-6

15. Mangel JC, Alfaro-Shigueto J, Witt MJ, Hodgson DJ, Godley BJ. Using pingers to reduce bycatch of small cetaceans in Peru’s small-scale driftnet fishery. Oryx. 2013; 47(4): 595–606. https://doi.org/10.1017/S0030605312000658

16. Carretta JV, Barlow J, Enriquez L. Acoustic pingers eliminate beaked whale bycatch in a gill net fishery. Mar Mamm Sci. 2008; 24(4): 956–961. https://doi.org/10.1111/j.1748-7692.2008.00218.x

17. Barlow J, Cameron GA. Field experiments show that acoustic pingers reduce marine mammal bycatch in the field. Mar Mamm Sci. 2003; 19(2): 265–283. https://doi.org/10.1111/j.1748-7692.2003.tb01108.x

18. Kraus SD, Read AJ, Solow A, Baldwin K, Spradlin T, Anderson E, et al. Acoustic alarms reduce porpoise mortality. Nature. 1997; 388: 525

19. Erbe C, Wintner S, Dudley SFJ, Plön S. Revisiting acoustic deterrence devices: Long-term bycatch data from South Africa’s bather protection nets. Proc Mtgs Acoust. 2016; 27: 010025. https://doi.org/10.1121/2.0000306

20. Soto AB, Cagnazzi D, Everingham Y, Parra GJ, Noad M, Marsh H. Acoustic alarms elicit only subtle responses in the behaviour of tropical coastal dolphins in Queensland, Australia. Endanger Species Res. 2013; 20(3): 271–282. https://doi.org/10.3354/esr00495

21. Nowacek DP, Johnson MP, Tyack PL. North Atlantic right whales (*Eubalaena glacialis*) ignore ships but respond to alerting stimuli. Proceedings of the Royal Society B: Biological Sciences. 2004; 271(1536): 227–231. https://doi.org/10.1098/rspb.2003.2570

22. Todd S, Lien J, Verhulst A. Orientation of humpback whales (Megaptera novaeangliae) and minke whales (Balaenoptera acutorostrata) to acoustic alarm devices designed to reduce entrapments in fishing gear. In: Thomas J, et al., editors. Marine Mammal Sensory Systems. Plenum Press: New York; 1992

23. Lagerquist B, Winsor M, Mate B. Testing the effectiveness of an acoustic deterrent for gray whales along the Oregon coast. Final Scientific Report. Oregon State University Marine Mammal Institute. U.S. Department of Energy; 2012 December. Available from: https://ir.library.oregonstate.edu/downloads/05741s112

24. Pirotta V, Slip D, Jonsen ID, Peddemors VM, Cato DH, Ross G, et al. Migrating humpback whales show no detectable response to whale alarms off Sydney, Australia. Endanger Species Res. 2016; 29(3): 201–209. https://doi.org/10.3354/esr00712

25. How J, Coughran DK, Smith JN, Harrison J, McMath J, Hebiton B, et al. Effectiveness of mitigation measures to reduce interactions between commercial fishing gear and whales. FRDC Project No 2013/03. In: Fisheries Research Report 267. Science; 21 October 2015: 635–645.

26. Harcourt R, Pirotta V, Heller G, Peddemors V, Slip D. A whale alarm fails to deter migrating humpback whales: An empirical test. Endanger Species Res. 2014; 25(1): 35–42. https://doi.org/10.3354/esr00614

27. Lien J, Barney W, Todd S, Seton R, Guzzwell J. Effects of Adding Sounds to Cod Traps on the Probability of Collisions by Humpback Whales. In: Thomas J, et al., editors. Marine Mammal Sensory Systems. Plenum Press: New York; 1992

28. Nowacek DP, Thorne LH, Johnston DW, Tyack PL. Responses of cetaceans to anthropogenic noise. Mamm Rev. 2007; 37(2): 81–115

29. Dunlop RA, Noad MJ, Cato DH, Kniest E, Miller PJO, Smith JN, et al. Multivariate analysis of behavioural response experiments in humpback whales (*Megaptera novaeangliae*). J Exp Biol. 2013; 216(5): 759–770. https://doi.org/10.1242/jeb.071498

30. Fumunda. ‘Pingers’ show promise to keep whale away from nets. Alaska Journal of Commerce.14 June 2012. Available from: http://www.alaskajournal.com/business-and-finance/2012-06-14/pingers-show-promise-keep-whales-away-nets

31. Welch L. Rebates help Alaska fishermen afford whale-repelling pingers. Anchorage Daily News. 31 May 2016. Available from: https://www.adn.com/business/article/rebates-help-alaska-fishermen-afford-whale-repelling-pingers/2016/05/07/

32. Taylor VJ, Johnston DW, Verboom WC. Acoustic harassment device (AHD) use in the aquaculture industry and implications for marine mammals. Proc. Symposium on Bio-sonar and Bioacoustics, Loughborough University, UK; 1997. Available from https://www.researchgate.net/profile/Willem_Verboom2/publication/279424748_Acoustic_Harassment_Device_AHD_use_in_the_aquaculture_industry_and_implications_for_marine_mammals/links/559266d108aed6ec4bf87c80/Acoustic-Harassment-Device-AHD-use-in-the-aquacult

33. McGarry T, Boisseau O, Stephenson S, Compton R. Understanding the Effectiveness of Acoustic Deterrent Devices (ADDs) on Minke Whale (*Balaenoptera acutorostrata*), a Low Frequency Cetacean. Offshore Renewables Joint Industry Programme (ORJIP) Project 4, Phase 2. Carbon Trust Offshore Renewables Joint Industry; 2017 December. Available from: https://www.carbontrust.com/media/675268/offshore-renewables-joint-industry-programme.pdf

34. Brandt MJ, Höschle C, Diederichs A, Betke K, Matuschek R, Nehls G. Seal scarers as a tool to deter harbour porpoises from offshore construction sites. Mar Ecol Prog Ser. 2013; 475: 291–302. https://doi.org/10.3354/meps10100

35. Morton AB, Symonds HK. Displacement of *Orcinus orca* (L.) by high amplitude sound in British Columbia, Canada. ICES J Mar Sci. 2002; 59(1): 71–80. https://doi.org/10.1006/jmsc.2001.1136

36. Morton A. Occurrence, photo-identification and prey of pacific white-sided dolphins (*Lagenorhyncus obliquidens*) in the Broughton Archipelago, Canada 1984-1998. Mar Mamm Sci. 2000; 16(1): 80-93

37. Pike DG, Paxton CG, Gunnlaugsson T, Víkingsson GA. (2009). Trends in the distribution and abundance of cetaceans from aerial surveys in Icelandic coastal waters, 1986-2001. NAMMCO Sci Publ. 2009; 7: 1986–2001. https://doi.org/10.7557/3.2710

38. Magnúsdóttir EE, Rasmussen MH, Lammers MO, Svavarsson J. Humpback whale songs during winter in subarctic waters. Polar Biol. 2013; 37(3): 427–433. https://doi.org/10.1007/s00300-014-1448-3

39. Sigurjónsson J, Víkingsson GA. Seasonal abundance of the estimated food consumption by cetaceans in Icelandic and adjacent waters. J Northwest Atl Fish Sci. 1997; 22: 271–287. https://doi.org/10.2960/J.v22.a20

40. Houser DS, Helweg DA, Moore PWB. A Bandpass filter-bank model of auditory sensitivity in the humpback whale. Aquat Mamm. 2001; 27(2): 82–91. https://doi.org/10.1103/PhysRevLett.89.180402

41. Todd SK. Acoustical properties of fishing gear; possible relationships to baleen whale entrapment. B.Sc.Hons Thesis, Memorial University of Newfoundland. 1991. Available from: https://research.library.mun.ca/5875/

42. Statistics Iceland. The fishing fleet by region and type of vessels 1999-2018 [cited 2019 Aug 15]. Available from: https://px.hagstofa.is/pxen/pxweb/en/Atvinnuvegir/Atvinnuvegir_sjavarutvegur_skip/SJA05001.px/table/tableViewLayout1/?rxid=23a75af5-f604-47a5-b3e9-62b122a29b55

43. Hafrannsóknastofnun. Fisheries overview. Hafrannsóknastofun, Reykjavik; 2018 June 13. Available from: https://www.hafogvatn.is/static/files/Veidiradgjof/2018/fishoverview_2018.pdf

44. Young, M. Marine animal entanglements in mussel aquaculture gear: Documented cases from mussel farming regions of the world including first-hand accounts from Iceland. MRM Thesis, University Center of the Westfjords. 2015. Available from: https://skemman.is/handle/1946/22522

45. Government of Iceland. Aquaculture. n.d. Ministry of Industries and Innovation. [cited 16 July 2019]. Available from: https://www.government.is/topics/business-and-industry/fisheries-in-iceland/aquaculture/

46. Basran CJ, Bertulli CG, Cecchetti A, Rasmussen MH, Whittaker M, Robbins J. First estimates of entanglement rate of humpback whales Megaptera novaeangliae observed in coastal Icelandic waters. Endanger Species Res. 2019; 38: 67–77. https://doi.org/10.3354/ESR00936

47. Basran C. Scar-based analysis and eyewitness accounts of entanglement of humpback whales (Megaptera novaeangliae) in fishing gear in Iceland. MRM Thesis, University Center of the Westfjords. 2014. Available from: https://skemman.is/handle/1946/19615

48. Víkingsson GA, Olafsdottir D, Gunnlaugsson Th. Iceland: Progress report on cetacean research, April 2003 to April 2004, with statistical data for the calendar year 2003. Marine Research Institute, Reykjavik. 2004. Document SC/ 56/ Prog. Rep. Available from: http://nammco.wpengine.com/wp-content/uploads/2016/08/Annual-Report-2005.pdf

49. Víkingsson GA, Olafsdottir D. Iceland: Progress report on cetacean research, April 2002 to March 2003, with statistical data for the calendar year 2002. Marine Research Institute, Reykjavik. 2003. Document SC/55/Prog.Rep. Available from: http://nammco.wpengine.com/wp-content/uploads/2016/08/Annual-Report-2004.pdf

50. Einarsson N. From good to eat to good to watch: whale watching, adaptation and change in Icelandic fishing communities. Polar Res. 2009; 28: 129−138.

51. Stefánsson U, Thórdardóttir Th, Ólafsson J. Comparison of seasonal oxygen cycles and primary production in the Faxaflói region, southwest Iceland. Deep Sea Res. 1987; 34: 725−739.

52. Stefánsson U, Guðmundsson G. The freshwater regime of Faxaflói, southwest Iceland, and its relationship to meteorological variables. Estuar Coast Mar Sci. 1978; 6: 535−551

53. Carlson, C. A Review of Whale Watch Guidelines and Regulations Around the World. IWC. 2009; 2008: 1–149

54. Southall B L, DeRuiter SL, Friedlaender A, Stimpert AK, Goldbogen JA, Hazen E, et al. Behavioral responses of individual blue whales (*Balaenoptera musculus*) to mid-frequency military sonar. J Exp Biol. 2019; 222(5): jeb190637. https://doi.org/10.1242/jeb.190637

55. Zuur AF, Ieno EN, Walker NJ, Saveliev AA, Smith GM. Mixed Effects Models and Extensions in Ecology with R. Springer Science+Business Media; 2009.

56. Pinheiro J, Bates D, DebRoy S, Sarkar D, R Core Team. _nlme: linear and nonlinear mixed effects models. R package version 3.1-117. 2014. Available from: http://CRAN.R-project.org/package=nlme

57. Hothorn T, Bretz F, Westfall P. Simultaneous Inferencein General Parametric Models. Biom J. 2008; 50(3): 346–363

58. Bates D, Maechler M, Bolker B, Walker S. Fitting Linear Mixed-Effects Models Using lme4. J Stat Softw. 2015; 67(1): 1–48. doi:10.18637/jss.v067.i01

59. Sivle LD, Kvadsheim PH, Curé C, Isojunno S, Wensveen PJ, Lam FPA, et al. Severity of expert-identified behavioural responses of humpback whale, minke whale, and northern bottlenose whale to naval sonar. Aquat Mamm. 2015; 41(4): 469–502. https://doi.org/10.1578/AM.41.4.2015.469

60. Blair HB, Merchant ND, Friedlaender AS, Wiley DN, Parks SE. Evidence for ship noise impacts on humpback whale foraging behaviour. Biol Lett. 2016; 12(8): 20160005

61. Ovide BG. Using tag data to assess behaviour, vocal sounds, boat noise and potential effects on humpback whales (Megaptera novaeangliae) in response to whale watching boates in Skjálfandi Bay (Húsavík), Iceland. MRM Thesis, University Center of the Westfjords. 2017. Available from: https://skemman.is/handle/1946/28667

62. Johnson JH, Wolman AA. The Humpback Whale, *Megaptera novaeangliae*. Mar Ecol Prog Ser. 1984; 46(4): 30–37.

63. Whalberg M, Westerberg H. Hearing in fish and their reactions to sounds from offshore wind farms. Mar Ecol Prog Ser. 2005; 288: 295–309.

64. Friedlaender AS, Tyson RB, Stimpert AK, Read AJ, Nowacek DP. Extreme diel variation in the feeding behavior of humpback whales along the western Antarctic Peninsula during autumn. Mar Ecol Prog Ser. 2013; 494: 281–289. https://doi.org/10.3354/meps10541

65. Perkins JS, Beamish PC. Net Entanglements of Baleen Whales in the Inshore Fishery of Newfoundland. J Fish Res Board Can. 1979; 36(5): 521–528.

66. Butterworth A, Clegg I, Bass C. Untangled – Marine debris: a global picture of the impact on animal welfare and of animal-focused solutions. London: World Society for the Protection of Animals; 2012. Available from: https://www.researchgate.net/publication/263444260_Marine_debris_a_global_picture_of_the_impact_on_animal_welfare_and_of_animal-focused_solutions

67. Boye TK, Simon M, Madsen PT. Habitat use of humpback whales in Godthaabsfjord, West Greenland, with implications for commercial exploitation. J Mar Biol Assoc UK. 2010; 90(8): 1529–1538

68. Schaffar A, Madon B, Garrigue C, Canstantine R. Behavioural effects of whale-watching activities on an Endangered population of humpback whales wintering in New Caledonia. Endang Species Res. 2013; 19: 245–254

69. Tubelli AA, Zosuls A, Ketten DR, Yamato M, Mountain DC. A prediction of the minke whale (*Balaenoptera acutorostrata*) middle-ear transfer function. J Acoust Soc Am. 2012;132(5):3263–3272. doi:10.1121/1.4756950

70. Baleen whale responses to a high frequency active pinger: Implications for upper frequency hearing limits. J Acoust Soc Am. 2016; 140: 3412. https://doi.org/10.1121/1.4970965

